# MetaMap, an interactive webtool for the exploration of metatranscriptomic reads in human disease-related RNA-seq data

**DOI:** 10.1101/425439

**Authors:** LM Simon, G Tsitsiridis, P Angerer, FJ Theis

## Abstract

**Motivation:** The MetaMap resource contains metatranscriptomic expression data from screening >17,000 RNA-seq samples from >400 archived human disease-related studies for viral and microbial reads, so-called “metafeatures”. However, navigating this set of large and heterogeneous data is challenging, especially for researchers without bioinformatic expertise. Therefore, a user-friendly interface is needed that allows users to visualize and statistically analyse the data.

**Results:** We developed an interactive frontend to facilitate the exploration of the MetaMap resource. The webtool allows users to query the resource by searching study abstracts for keywords or browsing expression patterns for specific metafeatures. Moreover, users can manually define sample groupings or use the existing annotation for downstream analysis. The web tool provides a large variety of analyses and visualizations including dimension reduction, differential abundance analysis and Krona visualizations. The MetaMap webtool represents a valuable resource for hypothesis generation regarding the impact of the microbiome in human disease.

**Availability:** The presented web tool can be accessed at https://github.com/theislab/MetaMap

## Introduction

An increasing number of studies investigates the role of the human microbiome in both health and disease(Young, 2017). RNA sequencing technology (RNA-seq) is commonly used to profile human gene expression patterns under defined conditions, including human disease-related contexts, but its generic nature also enables the detection of non-human RNA, including microbial and viral transcripts(Westermann et al., 2012).

The MetaMap resource(Simon et al., 2018) harbours the results from screening close to 150 terabytes of raw, publicly available RNA-seq data from the Sequencing Read Archive (SRA)(Leinonen et al., 2011) for non-human RNA. This big data resource contains the detected expression levels of microbial and viral species, so-called “metafeatures”, across >17,000 samples from >400 studies relevant to human disease and represents a valuable tool for the hypothesis generation toward the role of the microbiome in human disease.

Due to both volume and complexity of the data as well as availability only in matrix form, users without bioinformatic expertise may find it challenging to explore this resource. A number of previous studies have shown that interactive webtools enable easy exploration of large and complex molecular data sets(Sun et al., 2018; Dumas et al., 2016; Ono et al., 2017). Therefore, we developed the user-friendly, interactive MetaMap webtool. Our webtool provides both analyses and visualizations of the database including 1) dimension reduction, 2) diversity analysis, 3) differential metafeature abundance analysis, 4) metafeaturemetafeature correlation analysis 5) metafeature composition plots, 6) phylogenetic sankey diagrams and 7) Krona visualizations(Ondov et al., 2011).

To share user sessions across brower instances, the MetaMap webtool provides a “bookmarking” feature with permanent reference URL. The presented webtool produces downloadable publication-ready customizable graphics. Furthemore, underlying expression data including sample annotation are made easily accessible and available for download. The R code used to generate this resource is freely available allowing easy extension of the webtool to related data resources.

## Methods

The MetaMap webtool was implemented using the R statistical software(R Core Team, 2013) in conjunction with the shiny package(RStudio, Inc, 2013) for building interactive web applications. The R phyloseq package(McMurdie and Holmes, 2013) is used heavily for visualization and analyses. Underlying metafeature expression data was downloaded from the GigaScience database(Simon et al.). Additional information from the metaSRA database(Bernstein et al., 2017) was integrated to improve sample annotation. The source code is publicly available at: https://github.com/theislab/MetaMap-web.

## Using the MetaMap webtool

To start exploring the data, the user needs to select a study of interest. This can be done by either 1) querying the study abstracts for a specific keyword, such as “nasopharyngeal carcinoma” or 2) browsing the expression patterns of metafeatures across all studies (Fig. 1A). After having chosen a single study of interest the user can select existing sample annotation from the SRA to define sample groupings. Alternatively, users can manually define sample groupings or restrict analysis to a given set of samples or metafeatures using the “Customize data” tab.

**Figure 1.**
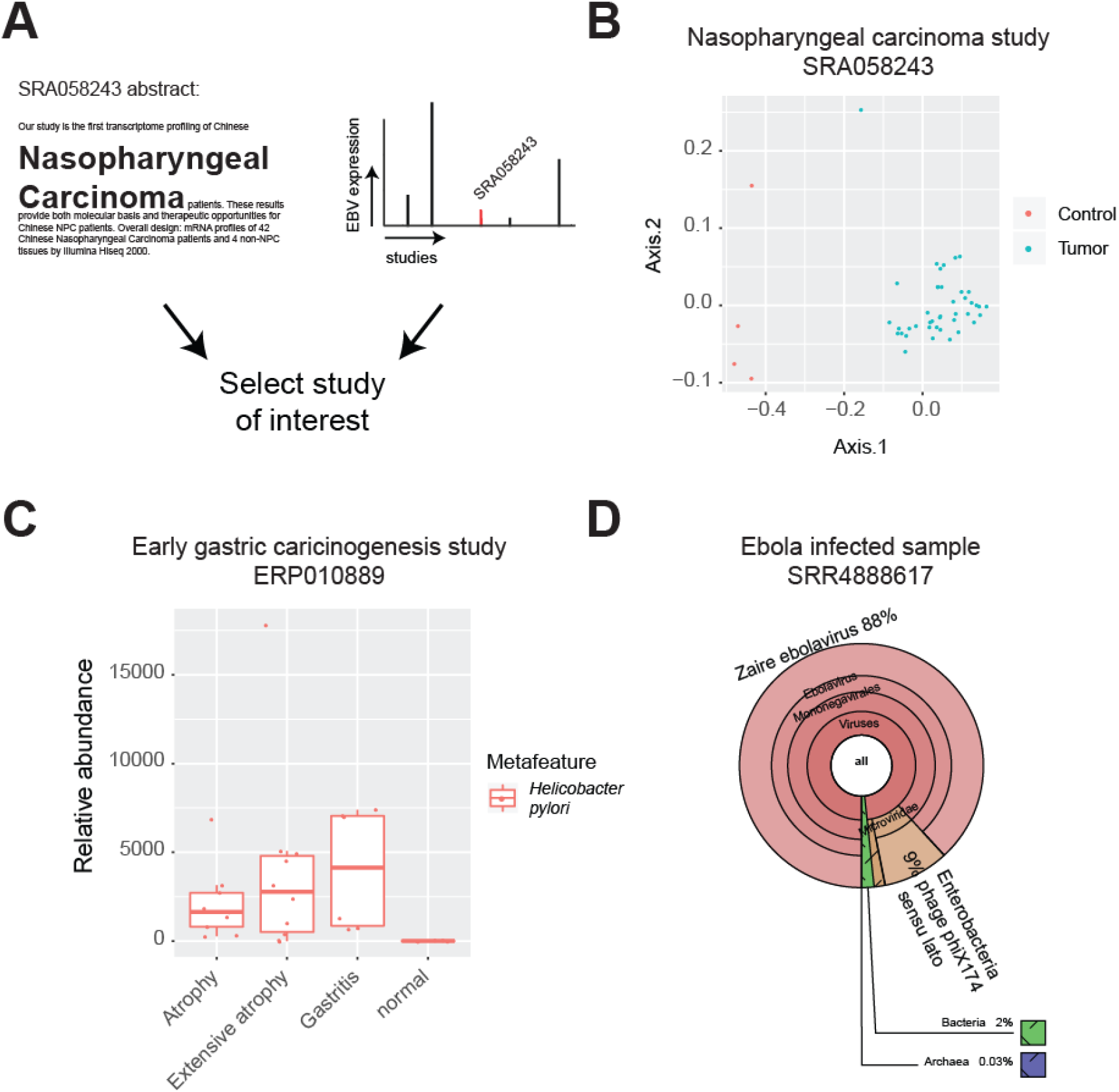
The MetaMap webtool provides interactive exploration of metatranscriptomic expression patterns across hundreds of studies relevant to human disease. (A) The user can select a study of interest by searching the SRA abstracts for keywords or by exploring the expression patterns of specific metafeatures. (B) MDS plot derived from metafeature expression data reveals clustering by cancer status. X and Y axes correspond to the MDS coordinates one and two. Samples are colored by tissue origin. (C) Differential metafeature abundance plot reveals differential expression of *Helicobacter pylori* across patient and control samples from SRA project ERP010889. Y axis represents metafeature abundance levels for *Helicobacter pylori* in reads per million total sequenced reads. (D) Krona visualization of Ebola infected sample illustrates high levels of the *Zaire ebolavirus* metafeature.

Next, the user can perform a number of analyses and generate various visualizations within the selected study. For example, dimension reduction of metafeature abundances using multidimensional scaling (MDS)(Gower et al., 1996) distinguished tumor samples from controls in a nasopharyngeal carcinoma study (SRA058243), implicating differential expression of underlying metafeatures in the disease (Fig. 1B, click here and see for yourself).

In another study (ERP010889) RNA-seq profiles were generated from gastric corpus biopsies to study early gastric carcinogenesis. The MetaMap webtool revealed that *Helicobacter pylori* was detected in “(Extensive) Atrophy” and “Gastritis” samples but not in healthy controls (Fig. 1C, click here and see for yourself). This finding explains using the MetaMap resource and demonstrates the value of the MetaMap webtool as well-known metafeature-disease associations can easily be (re-)discovered(Ebule et al., 2013).

A third exemplary study (SRP092544) investigated transcriptome data for peripheral blood taken from Ebola virus infected patients. Krona visualization of metafeature expression derived from a patient sample showed 88% of metafeature reads classified as *Zaire ebolavirus* (Fig 1D, click here and see for yourself).

Overall these three examples showcase the value of the MetaMap webtool, but do no exhaust the potential for further findings. In conclusion, the MetaMap webtool represents a valuable, user-friendly resource for the hypothesis generation toward the role of the microbiome in human disease.

